# A High Quality Assembly of the Nile Tilapia (*Oreochromis niloticus*) Genome Reveals the Structure of Two Sex Determination Regions

**DOI:** 10.1101/099564

**Authors:** Matthew A Conte, William J Gammerdinger, Kerry L Bartie, David J Penman, Thomas D Kocher

## Abstract

We report a high-quality assembly of the tilapia genome, a perciform fish important in aquaculture around the world. A homozygous clonal XX female Nile tilapia (*Oreochromis niloticus*) was sequenced to 44X coverage using Pacific Biosciences (PacBio) SMRT sequencing. Dozens of candidate *de novo* assemblies were generated and an optimal assembly (contig NG50 of 3.3Mbp) was selected using principal component analysis of likelihood scores calculated from several paired-end sequencing libraries. Comparison of the new assembly to the previous *O. niloticus* genome assembly reveals that recently duplicated portions of the genome are now well represented. The overall number genes in the new assembly increased by 27.3%, including a 67% increase in pseudogenes. The new tilapia genome assembly correctly represents two recent *vasa* gene duplication events that have been verified with BAC sequencing. At total of 146Mbp of additional transposable element sequence are now assembled, a large proportion of which are recent insertions. Large centromeric satellite repeats are assembled and annotated in cichlid fish for the first time. Finally, the new assembly identifies the long-range structure of both an ~9Mbp XY sex-determination region on LG1 in *O. niloticus*, and a ~50Mbp WZ sex-determination region on LG3 in the related species *O. aureus.* This study highlights the use of long read sequencing to correctly assemble recent duplications and to characterize repeat-filled regions of the genome.

## Introduction

Aquaculture plays an increasingly important role in providing sustainable seafood products and has significantly outpaced capture fisheries in the past several decades (1). Tilapias are among the most important farmed fishes, and tilapia production continues to expand exponentially across the globe (2). An important aspect of commercial production is the control of sexual differentiation. Males grow to market-size earlier than females. Females also start to reproduce at a smaller size, filling production ponds with small fish (3). It is therefore advantageous to grow-out only male fish. At one time, all-male populations were produced through interspecific crosses (4), but the strains supporting this technology have been lost or contaminated. Currently, the standard way of achieving all male or nearly all male tilapia populations is via hormonal masculinization (3,5). A reliable way of producing genetically all-male tilapia would allow the replacement of hormonal masculinization, which is banned in several major producing countries (although not enforced in most cases). It is therefore important to understand the genetic basis of sex determination in current aquaculture stocks.

Sex determination in tilapias is largely genetic, although environmental factors also play a role (6–8). In Nile tilapia (*Oreochromis niloticus*), distinct XY sex determining loci have been identified on both LGs 1 and 23 (9,10). The closely related blue tilapia (*O. aureus*) segregates both an XY locus on LG1, and an epistatically dominant ZW locus on LG3 (11). Additional sex determining loci have been identified on LGs 5, 7, 13, 18 and 20 in closely related species of East African cichlid (12–14). As a group, tilapias and related species of other cichlid fishes are a promising model system for understanding the gene network controlling sex determination in vertebrates.

Work to identify the genes underlying each of these sex determiners has been hampered by the incomplete nature of previous draft genome assemblies, and by the discovery that many of these sex determiners are located in large blocks of highly differentiated, and sometimes repetitive sequence. To date, the molecular genetic basis for sex determination in cichlids has been determined for only the LG23 XY locus in *O. niloticus* (15).

Although several draft genome sequences are available for cichlids, these are mostly based on short Illumina sequencing reads (16). Recently duplicated and highly repetitive sequences are typically collapsed in these assemblies (17). Indeed some of the most interesting and perhaps evolutionarily important regions of the genome may be the most difficult to assemble accurately. Recently duplicated regions are notoriously difficult to assemble due to their repeat length and high sequence identity (18). The repetitive “dark-matter” part of the genome is vastly underrepresented in the majority of current genome assemblies (19). Attempts to assemble these regions using only short read sequencing are futile (20). Only long sequencing reads will produce more contiguous and complete assemblies of complex vertebrate genomes (21–25). The importance of such high quality assemblies for downstream applications cannot be overemphasized.

Here we report a new assembly of the tilapia genome from long PacBio sequence reads. This assembly contains much of the missing sequence from previous assemblies, and is among the most contiguous vertebrate genome assemblies to date. We use this new assembly to further characterize the tilapia sex determining loci previously identified on LGs 1 and 3.

## Results

### Assembly Overview

A homozygous clonal XX female tilapia individual (26) was chosen for genome sequencing. The individual was sequenced to 44X coverage using PacBio sequencing of 63 SMRT cells using the P6-C4 chemistry. This yielded 5,085,371 reads with a mean subread length of 8,747bp and N50 read length of 11,366bp.

An overview of the assembly process is outlined in Fig 1. To summarize, 37 candidate *de novo* assemblies were generated using both the FALCON (22) and Canu (27) genome assembly packages. Multiple parameters were adjusted for both algorithms to tune the assemblies. The error correction steps of both algorithms include parameters that control alignment seed length, read length, overlap length and error rates (see Methods).

**Fig 1.**
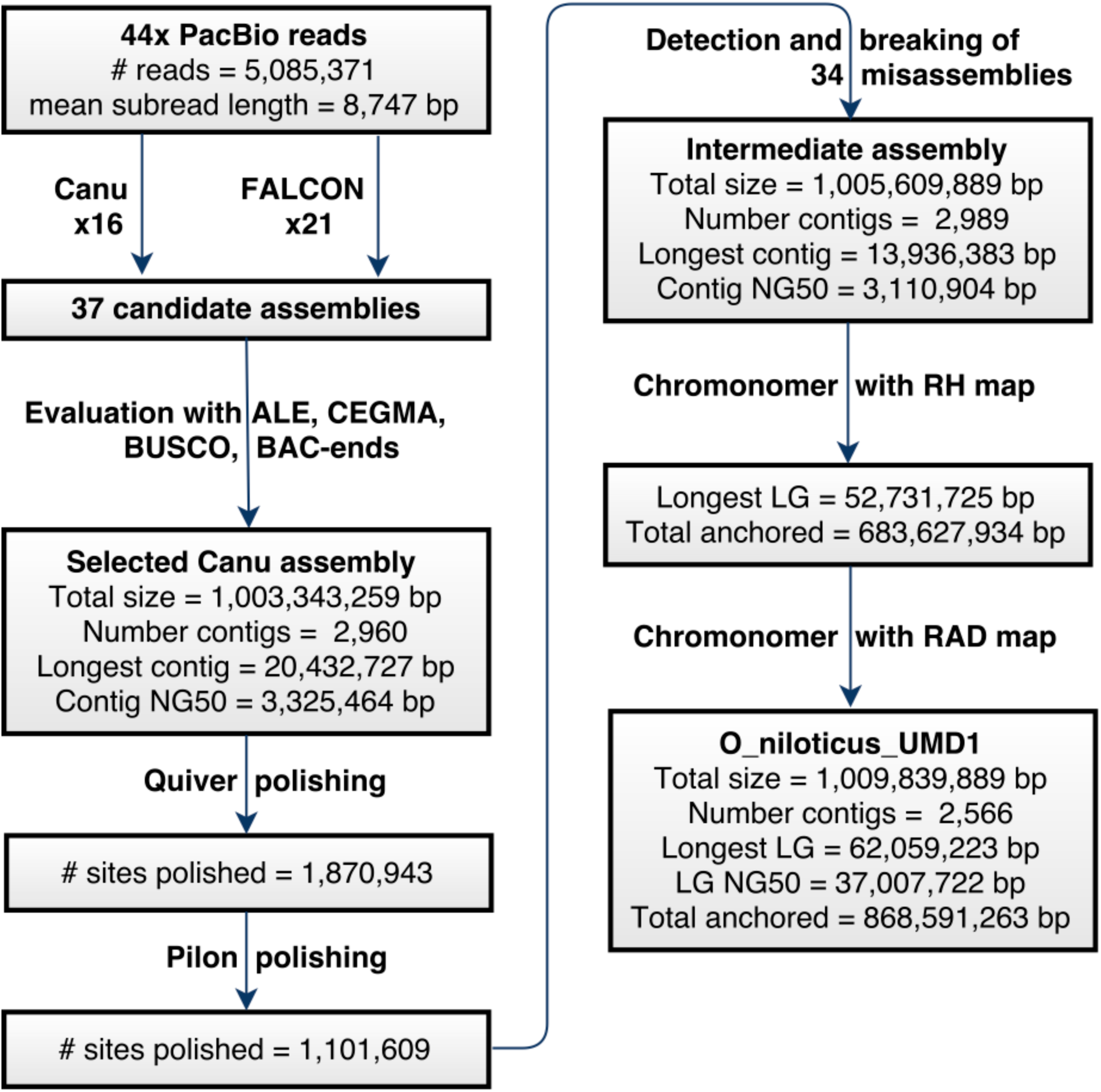
Assembly overview. Flowchart detailing the processing of the raw 44X PacBio sequencing reads, producing candidate assemblies, polishing, breaking, and final assembly anchoring. Metrics are provided at each step.

### Evaluating Assemblies

The 37 candidate assemblies were evaluated using a number of different metrics, techniques and complementary datasets. First, each of the candidate assemblies was evaluated using ALE assembly likelihood estimates (28) (which integrated read quality, mate-pair orientation, insert size, coverage and *k*-mer frequencies) based on alignment of the reads from four separate Illumina libraries and of the 44X PacBio dataset (see Methods). Candidate assemblies were also evaluated for completeness using CEGMA (29) and BUSCO (30) core gene sets, as well as by aligning existing *O. niloticus* RefSeq (31) transcripts. A set of 193,027 BAC-end sequences (32) representing ~29X physical clone coverage were used to assess the longer range accuracy of candidate assemblies. Finally, both a physical radiation-hybrid (RH) map consisting of 1,256 markers (33) and a RAD-seq genetic map consisting of 3,802 markers (34) were used to estimate the number of misassemblies present in each of the candidate assemblies. The results of these analyses are provided in Table S1.

### Ranking Assemblies

No single candidate assembly ranked the highest for all of the evaluation metrics that were computed. Principal component analysis (PCA) was used to reduce the various assembly evaluation metrics and compare the candidate assemblies. S1 Fig shows that the Canu assemblies tend to cluster separately from the FALCON assemblies in the PCA space. The total assembly size and number of RefSeq exons mapped explained the largest amount of variance and were correlated. These two metrics did not seem like the most important metrics to base the evaluation upon since assembly parameters could be tuned to change the total size and the estimated genome size was 1.082Gbp (35).

The ALE likelihood scores explain the next largest proportion of the variance. The 37 candidate assemblies were ranked by overall ALE scores for each of the five sequencing libraries. An average of the ALE ranks was then calculated. The Canu assembly (#14) that was chosen as the best among the 37 candidate assemblies showed the best average ALE ranks. In addition, Canu assembly #14 had one of the best rates of properly mapped BAC-end sequences, and possessed among the fewest misassemblies as determined by conflicts with the RH and RAD map data (Table S1). These results suggest that Canu assembly #14 has the best long-range accuracy while maintaining comparable short-range accuracy.

### Polishing

A relatively small number of sequence errors remained in the intermediate unpolished Canu #14 assembly. To correct these errors, first the raw 44X PacBio reads were aligned to the Canu assembly and Quiver was used to polish the assembly at 1,870,943 sites (see Methods). Quiver corrected 1,739,112 (92.95%) insertions, 88,037 (4.71%) substitutions and 43,794 (2.34%) deletions. Next, four Illumina libraries, totaling 277x coverage, were aligned to the intermediate Quiver-polished assembly. Based upon these alignments, Pilon polished an additional 1,101,609 sites. Pilon corrected 1,087,107 (98.68%) insertions, 12,402 (1.13%) substitutions, and 2,100 (0.19%) deletions.

### Detection of Misassemblies

The polished intermediate assembly showed high accuracy at the level of individual bases and with respect to the placement of paired-end sequences from ~150kbp BACs (Table S1). However, 32 putative inter-chromosomal misassemblies were identified by alignment to the RH and RAD maps. The RH and RAD maps both identified 21 of these inter-chromosomal misassemblies. The RAD map identified an additional 8 putative misassemblies that were not identified using the RH map (the RH map had no markers aligning to these regions), while the RH map identified an additional 3 misassemblies that were not identified using the RAD map (likewise, the RAD map had no markers aligning to these regions). The regions around each putative misassembly were inspected using the genomic resources already mentioned. Each had a characteristic signature consisting of a high density of variants in the 44X PacBio read alignments, as well as low or zero physical coverage of the 40kbp insert Illumina mate-pair library. An example of these misassembly signatures is shown in S2 Fig.

Genome wide analysis of the intermediate assembly for each of these characteristic signatures detected 110 regions of high-density PacBio variants and 376 regions of low physical coverage in the 40kbp mate-pair library. 41 regions had both a high-density of PacBio variants and low physical 40kbp mate-pair coverage. Nine of these regions showed correct alignment to both maps and therefore were not included in the set of putative misassemblies. However, two of these regions were identified by the PacBio variants and low 40kbp physical coverage in regions where there were no markers in either the RH or RAD map and added to the 32 map-based misassemblies giving a total of 34 sites of likely misassembly. Table 1 provides a summary of the putative misassemblies that were identified by the maps and sequence alignment methods.

**Table 1.** Number of putative misassemblies identified by various methods.

| Evidence | Number of misassemblies detected |
| --- | --- |
| Both maps | 21 |
| RAD map only | 8 |
| RH map only | 3 |
| Both PacBio variants and 40kbp library | 16 |
| PacBio variants only | 2 |
| 40kbp library only | 16 |
| PacBio variants and 40kbp library, but no maps | 2 |

Analysis of the repetitive elements within these regions revealed that misassembly locations were enriched for highly repetitive interspersed and nested repeats. We examined the region ~75kbp on both sides of the likely misassembly breakpoints and found that 94.51% of these regions were classified as repetitive (see Methods). These regions were enriched for several TE families. Table 2 shows the enrichment of the most common repeats and TEs within the misassembly regions. In each of these cases, the mean length of these repeats was longer within the misassembly regions. Some of the same TE families that are abundant across the whole genome (e.g. DNA-TcMar-Tc1, LINE-L2, LINE-Rex-Babar) are also present in high frequency in the misassembly regions. However, some TE families that occurred in relatively low frequency across the whole genome (e.g. DNA-Sola, LTR-ERV1, RC-Helitron, and satellite repeats) are highly enriched in the misassembly regions.

**Table 2.** Repeats in putative misassembly regions compared to the whole genome.

| Repetitive element |  | O_niloticus_UMD1 genome wide |  |  |  | Within 34 misassembly regions |  |  |  | Enrichment |  |
| --- | --- | --- | --- | --- | --- | --- | --- | --- | --- | --- | --- |
| Order | Family | # | Total bp | % | Mean length | # | Total bp | % | Mean length | Ratio of genome/<br>missassem<br>regions | Δ<br>mean<br>length |
| **DNA** | Sola | 7,007 | 1,536,337 | 0.15% | 219.3 | 142 | 53,045 | 1.37% | 373.6 | 913.3% | 154.3 |
|  | TcMar-Tc1 | 156,588 | 46,394,192 | 4.60% | 296.3 | 846 | 354,606 | 9.18% | 419.2 | 199.6% | 122.9 |
|  | hAT | 36,441 | 10,103,158 | 1.00% | 277.2 | 289 | 99,016 | 2.56% | 342.6 | 256.0% | 65.4 |
|  | hAT-Ac | 49,528 | 17,626,929 | 1.75% | 355.9 | 445 | 191,054 | 4.95% | 429.3 | 282.9% | 73.4 |
|  | hAT-Charlie | 37,049 | 13,709,558 | 1.36% | 370.0 | 236 | 99,913 | 2.59% | 423.4 | 190.4% | 53.3 |
| **LINE** | L1 | 10,712 | 8,879,041 | 0.88% | 828.9 | 63 | 79,410 | 2.06% | 1,261 | 234.1% | 431.6 |
|  | L2 | 76,937 | 29,334,193 | 2.91% | 381.3 | 539 | 269,645 | 6.98% | 500.3 | 239.9% | 119.0 |
|  | Penelope | 28,509 | 7,214,522 | 0.71% | 253.1 | 258 | 70,627 | 1.83% | 273.7 | 257.7% | 20.7 |
|  | Rex-Babar | 38,996 | 19,208,630 | 1.90% | 492.6 | 386 | 199,730 | 5.17% | 517.4 | 272.1% | 24.9 |
| **LTR** | ERV1 | 10,756 | 6,450,995 | 0.64% | 599.8 | 112 | 97,055 | 2.51% | 866.6 | 392.2% | 266.8 |
|  | Gypsy | 29,201 | 13,615,743 | 1.35% | 466.3 | 274 | 172,231 | 4.46% | 628.6 | 330.4% | 162.3 |
| **RC** | Helitron | 3,882 | 2,685,111 | 0.27% | 691.7 | 216 | 187,805 | 4.86% | 869.5 | 1800.0% | 177.8 |
| **Unk.** | Unknown | 350,007 | 97,456,302 | 9.66% | 278.4 | 2,188 | 841,264 | 21.78% | 384.5 | 225.5% | 106.0 |
| **Satellite** | Satellite | 10,061 | 7,662,597 | 0.76% | 761.6 | 93 | 115,309 | 2.99% | 1,240 | 393.4% | 478.3 |
|  | Simple | 322,732 | 15,543,664 | 1.54% | 48.2 | 1,733 | 173,824 | 4.50% | 100.3 | 292.2% | 52.1 |

### Anchoring

Table 3 provides the anchored size of each LG, including gaps. The new O_niloticus_UMD1 assembly anchored 868.6Mbp of the total genome (86.9%), which is 211Mbp (32%) more than was anchored in the previous “Orenil1.1” assembly (657Mbp) (16). When gaps are not counted, the amount of anchored, non-gap, sequence is 864Mbp (86.4%) compared to 606Mbp (60.6%) in the previous Orenil1.1 assembly. LG3 is the largest anchored LG (68.6Mbp), which agrees with cytogenetic studies that show LG3 as the largest and most repetitive chromosome in the *O. niloticus* genome (33,36,37). Cytogenetic studies also indicate that LG7 is the second largest chromosome in the *O. niloticus* genome, and LG7 is the second largest anchored LG in the new “O_niloticus_UMD1” assembly.

**Table 3.** Size of each anchored linkage group for both the previous assembly, Orenil1.1 (16) and the new assembly (O_niloticus_UMD1).

| Linkage Group | Orenil1.1 | *O_niloticus_UMD1* | Difference (%) |
| --- | --- | --- | --- |
| LG1 | 31,194,787 | 38,372,991 | 7,178,204 (23.0%) |
| LG2 | 25,048,291 | 35,256,741 | 10,208,450 (40.8%) |
| LG3 | 19,325,363 | 68,550,753 | 49,225,390 (254.7%) |
| LG4 | 28,679,955 | 38,038,224 | 9,358,269 (32.6%) |
| LG5 | 37,389,089 | 34,628,617 | -2,760,472 (-7.4%) |
| LG6 | 36,725,243 | 44,571,662 | 7,846,419 (21.4%) |
| LG7 | 51,042,256 | 62,059,223 | 11,016,967 (21.6%) |
| LG8 | 29,447,820 | 30,802,437 | 1,354,617 (4.6%) |
| LG9 | 20,956,653 | 27,519,051 | 6,562,398 (31.3%) |
| LG10 | 17,092,887 | 32,426,571 | 15,333,684 (89.7%) |
| LG11 | 33,447,472 | 36,466,354 | 3,018,882 (9.0%) |
| LG12 | 34,679,706 | 41,232,431 | 6,552,725 (18.9%) |
| LG13 | 32,787,261 | 32,337,344 | -449,917 (-1.4%) |
| LG14 | 34,191,023 | 39,264,731 | 5,073,708 (14.8%) |
| LG15 | 26,684,556 | 36,154,882 | 9,470,326 (35.5%) |
| LG16 | 34,890,008 | 43,860,769 | 8,970,761 (25.7%) |
| LG17 | 31,749,960 | 40,919,683 | 9,169,723 (28.9%) |
| LG18 | 26,198,306 | 37,007,722 | 10,809,416 (41.3%) |
| LG19 | 27,159,252 | 31,245,232 | 4,085,980 (15.0%) |
| LG20 | 31,470,686 | 36,767,035 | 5,296,349 (16.8%) |
| LG22 | 26,410,405 | 37,011,614 | 10,601,209 (40.1%) |
| LG23 | 20,779,993 | 44,097,196 | 23,317,203 (112.2%) |
| **Total** | **657,350,972** | **868,591,263** | **211,240,291 (32.1%)** |
| **Total (minus gaps%)** | **606,480,097** | **864,361,263** | **257,881,166 (42.5%)** |

### Assembly Completeness

To determine the completeness of the new O_niloticus_UMD1 assembly, the assembly was compared against two established sets of core vertebrate gene sets. Table 4 shows the number of the 248 CEGMA and the 3,023 BUSCO conserved vertebrate genes that were identified in the new assembly. The number of conserved genes identified increased for both the CEGMA and BUSCO gene sets. The number of complete single-copy BUSCOs increased by 223 (10%), while the number of complete duplicated BUSCOs increased by 26 (59%). The number of missing BUSCOs decreased by 288 (67%) in the O_niloticus_UMD1 assembly compared to the Orenil1.1 assembly.

**Table 4.**
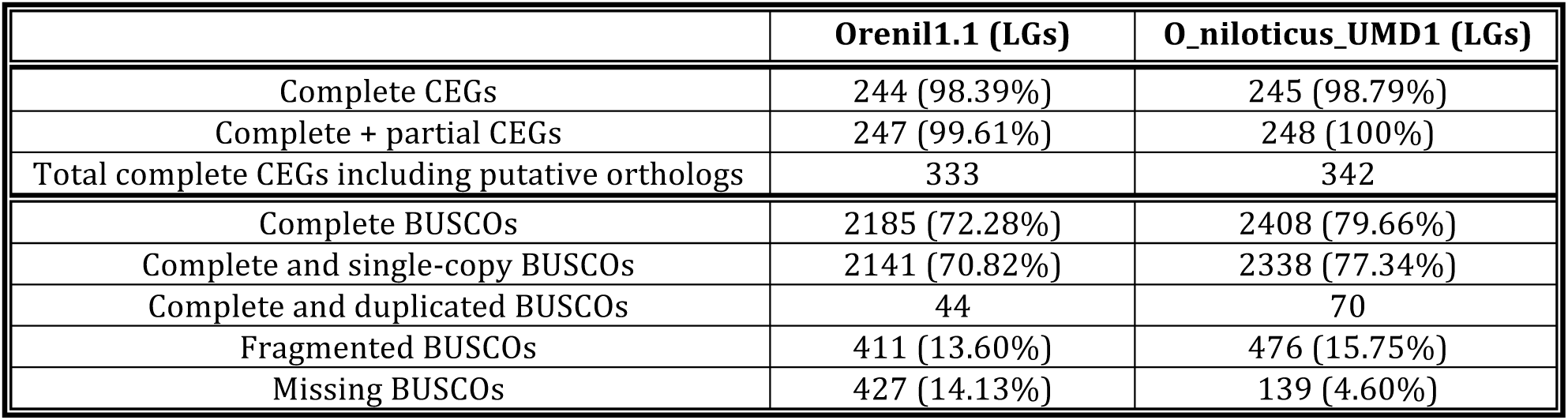
Genome completeness as measured by CEGMA and BUSCO.

|  | **Orenil1.1 (LGs)** | **O_niloticus_UMD1 (LGs)** |
| --- | --- | --- |
| Complete CEGs | 244 (98.39%) | 245 (98.79%) |
| Complete + partial CEGs | 247 (99.61%) | 248 (100%) |
| Total complete CEGs including putative orthologs | 333 | 342 |
| Complete BUSCOs | 2185 (72.28%) | 2408 (79.66%) |
| Complete and single-copy BUSCOs | 2141 (70.82%) | 2338 (77.34%) |
| Complete and duplicated BUSCOs | 44 | 70 |
| Fragmented BUSCOs | 411 (13.60%) | 476 (15.75%) |
| Missing BUSCOs | 427 (14.13%) | 139 (4.60%) |

### Annotation

The O_niloticus_UMD1 assembly was annotated using the NCBI RefSeq automated eukaryotic genome annotation pipeline. This same pipeline was previously used to annotate the Orenil1.1 assembly. A comparison of both genome assembly annotations is provided in Table 5. The O_niloticus_UMD1 assembly contains 8,238 more gene and pseudogene annotations than the Orenil1.1 assembly (27.3% increase). Similarly, the number of mRNA annotations increased markedly by 10,374 (21.7% increase). The number of partial mRNA annotations decreased from 3,050 to 393 (87.1% decrease). CDS annotations also increased overall (21.9%). The RefSeq annotation pipeline makes corrections to CDS annotations that contain premature stop-codons, frameshifts and internal gaps that would disrupt protein sequence coding. These corrections are based on transcriptome data and corrected 743 CDSs in O_niloticus_UMD1 compared to 817 previously for Orenil1.1 (9.1% decrease). The number of non-coding RNAs more than doubled in the O_niloticus_UMD1 assembly (115.5% increase).

**Table 5.**
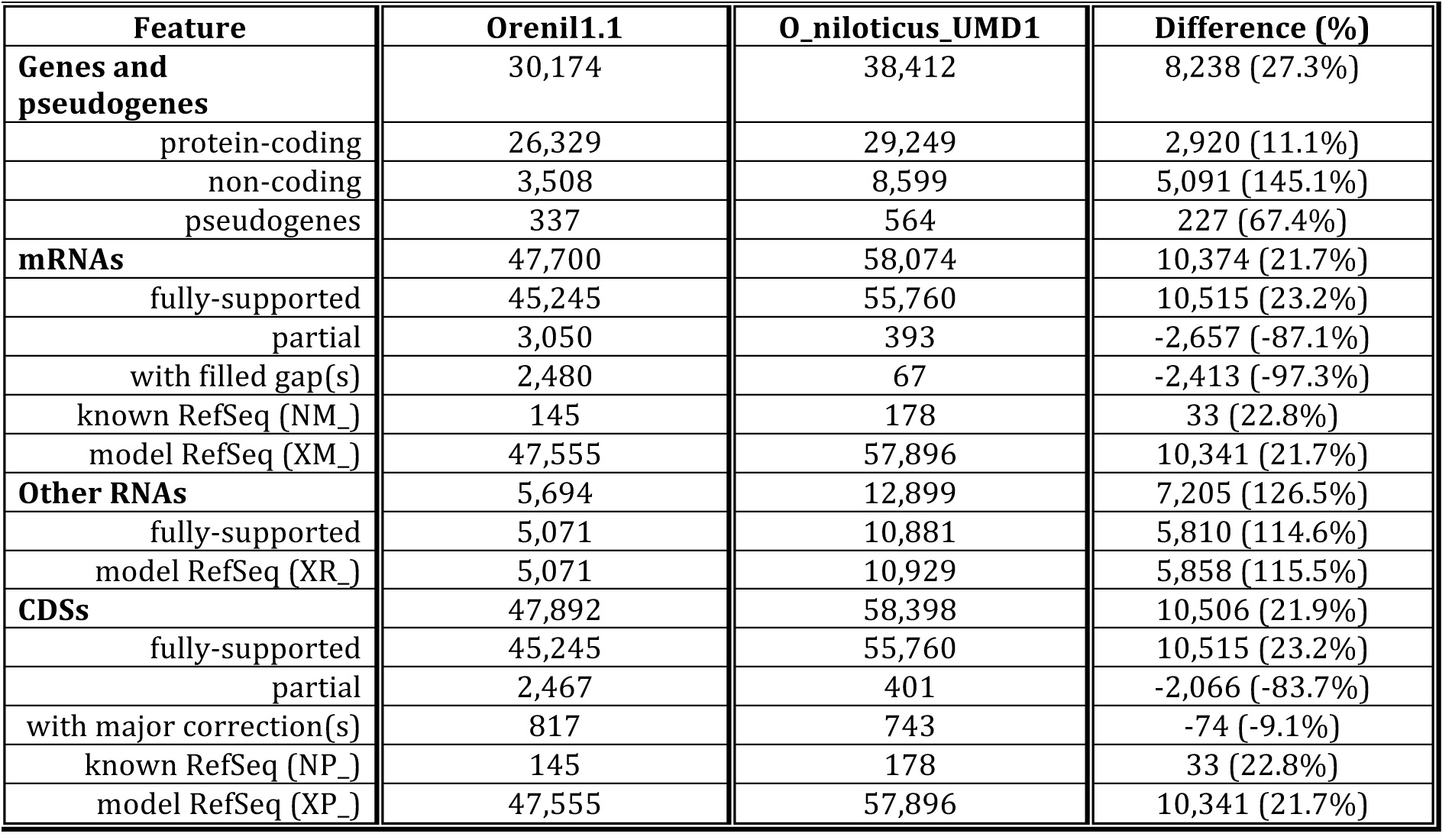
RefSeq annotation summary.

| Feature | Orenil1.1 | O_niloticus_UMD1 | Difference (%) |
| --- | --- | --- | --- |
| **Genes and pseudogenes** | 30,174 | 38,412 | 8,238 (27.3%) |
| protein-coding | 26,329 | 29,249 | 2,920 (11.1%) |
| non-coding | 3,508 | 8,599 | 5,091 (145.1%) |
| pseudogenes | 337 | 564 | 227 (67.4%) |
| **mRNAs** | 47,700 | 58,074 | 10,374 (21.7%) |
| fully-supported | 45,245 | 55,760 | 10,515 (23.2%) |
| partial | 3,050 | 393 | -2,657 (-87.1%) |
| with filled gap(s) | 2,480 | 67 | -2,413 (-97.3%) |
| known RefSeq (NM_) | 145 | 178 | 33 (22.8%) |
| model RefSeq (XM_) | 47,555 | 57,896 | 10,341 (21.7%) |
| **Other RNAs** | 5,694 | 12,899 | 7,205 (126.5%) |
| fully-supported | 5,071 | 10,881 | 5,810 (114.6%) |
| model RefSeq (XR_) | 5,071 | 10,929 | 5,858 (115.5%) |
| **CDSs** | 47,892 | 58,398 | 10,506 (21.9%) |
| fully-supported | 45,245 | 55,760 | 10,515 (23.2%) |
| partial | 2,467 | 401 | -2,066 (-83.7%) |
| with major correction(s) | 817 | 743 | -74 (-9.1%) |
| known RefSeq (NP_) | 145 | 178 | 33 (22.8%) |
| model RefSeq (XP_) | 47,555 | 57,896 | 10,341 (21.7%) |

### O_niloticus_UMD1 Assembly Summary

Table 6 provides summary statistics for the previous *O. niloticus* assembly (Orenil1.1), each intermediate of the new assembly, and our new final assembly (O_niloticus_UMD1). The O_niloticus_UMD1 assembly is more contiguous, with 45% fewer contigs than the number of scaffolds in Orenil1.1. The overall size of the O_niloticus_UMD1 assembly is 1.01Gbp compared to 927Mbp of Orenil1.1. The O_niloticus_UMD1 contains only 424 gaps that were introduced in the anchoring step. These anchoring gaps amount to 4.2Mbp (0.42%) due entirely to the arbitrary 10kbp gaps placed between anchored contigs. This compares to 111.5Mbp (12.04%) of gaps in Orenil1.1. Overall, 189.5Mbp of new sequence has been assembled in O_niloticus_UMD1 that was either previously in gaps or not assembled at all in Orenil1.1.

**Table 6.** Summary statistics for the various assemblies.

| Assembly | O_niloticus_wgs_v1<br>(scaffolds) | Orenil1.1<br>(anchored<br>to LGs) | *O. niloticus*<br>Canu<br>(contigs) | *O. niloticus*<br>Canu broken<br>(contigs) | O_niloticus_UMD1<br>(anchored<br>to LGs) |
| --- | --- | --- | --- | --- | --- |
| Number of<br>contigs/scaffolds | 5,900 | 5,677 | 2,960 | 2,989 | 2,566 |
| Total size | 927,725,912 | 927,679,487 | 1,003,343,259 | 1,005,609,889 | 1,009,839,889 |
| Longest<br>contig/scaffold/LG | 13,623,339 | 51,042,256 | 20,432,727 | 13,936,383 | 62,059,223 |
| Mean<br>contig/scaffold/LG<br>size | 157,242 | 29,879,589.6 | 338,967 | 336,437 | 37,672,228.8 |
| NG50<br>contig/scaffold/LG | 2,629,658 | 26,684,556 | 3,325,464 | 3,110,904 | 37,007,722 |
| % N | 12.04 | 12.03 | 0 | 0 | 0.42 |
| Number of gaps | 71,854 | 72,077 | 0 | 0 | 424 |
| Non gap bp | 816,139,901 | 816,140,124 | 1,003,343,259 | 1,005,609,889 | 1,005,610,312 |
| Total gap bp | 111,586,011 | 111,539,363 | 0 | 0 | 4,240,000 |

### Repeat Content

The TE and repeat portion of the genome is vastly under underrepresented in most genome assemblies (19). The use of long PacBio reads allowed for the assembly of more of the repetitive regions of the *O. niloticus* genome. 379Mbp (37.6%) of the total O_niloticus_UMD1 assembly was annotated as repetitive. Table 7 provides a summary of the repeat and TE families that were most abundant in the assembly. The new assembly includes an additional 146Mbp (14.6%) of repetitive sequence that was either hidden in gaps or not present at all in the previous assembly. The entire repeat catalog is provided in Table S2.

**Table 7.**
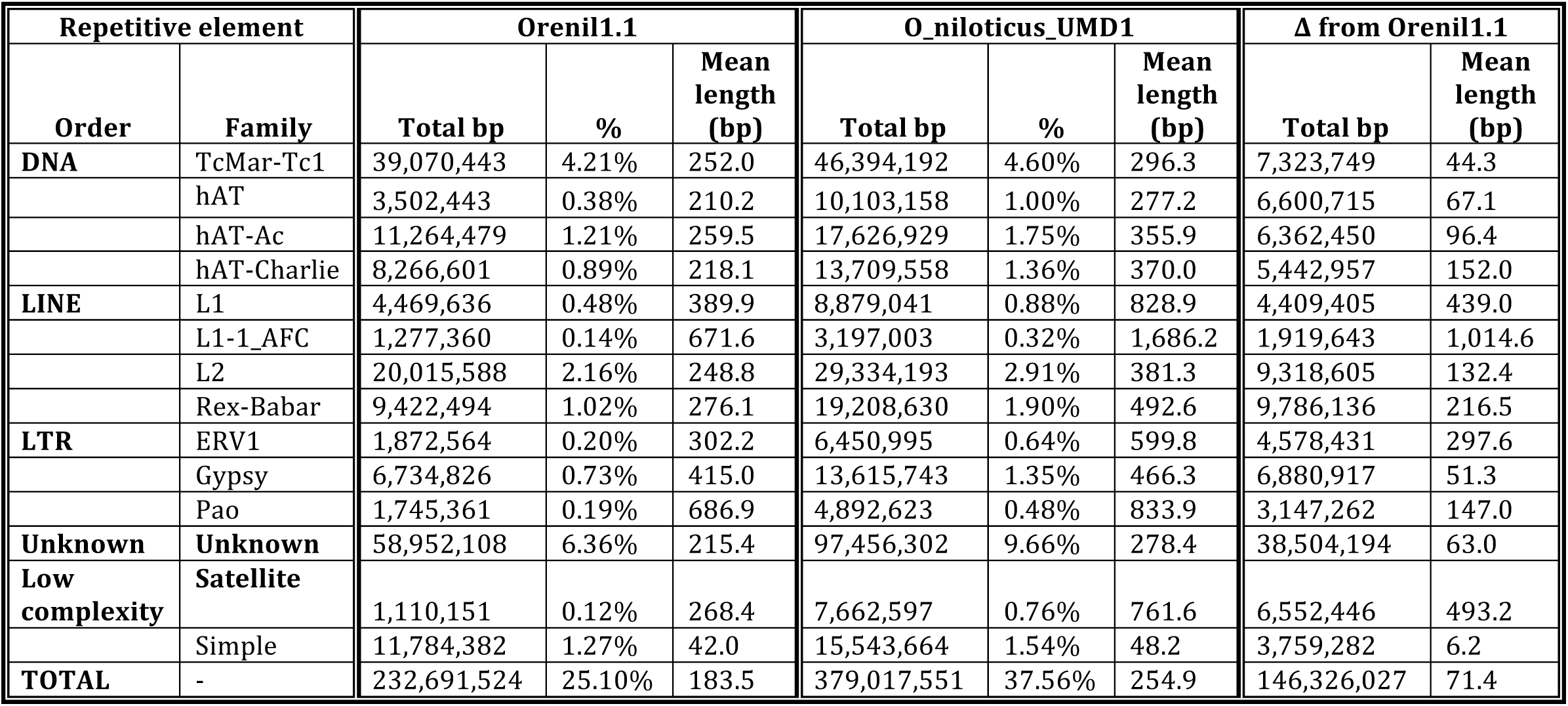
Summary of repeat families in the new assembly.

Fig 2 provides a comparison of the repeat landscape of the Orenil1.1 and O_niloticus_UMD1 assemblies. Most notably, recently inserted (~ < 5% Kimura divergence) TEs have been assembled in far greater number in the new O_niloticus_UMD1 assembly. The overall number of repetitive elements increased at all divergence levels (218,992 more elements, Table S2), with most at lower divergences (165,607 additional elements at < 5% Kimura divergence). The graph suggests that TE insertions less than 1% diverged are still underrepresented in the assembly.

**Fig 2.**
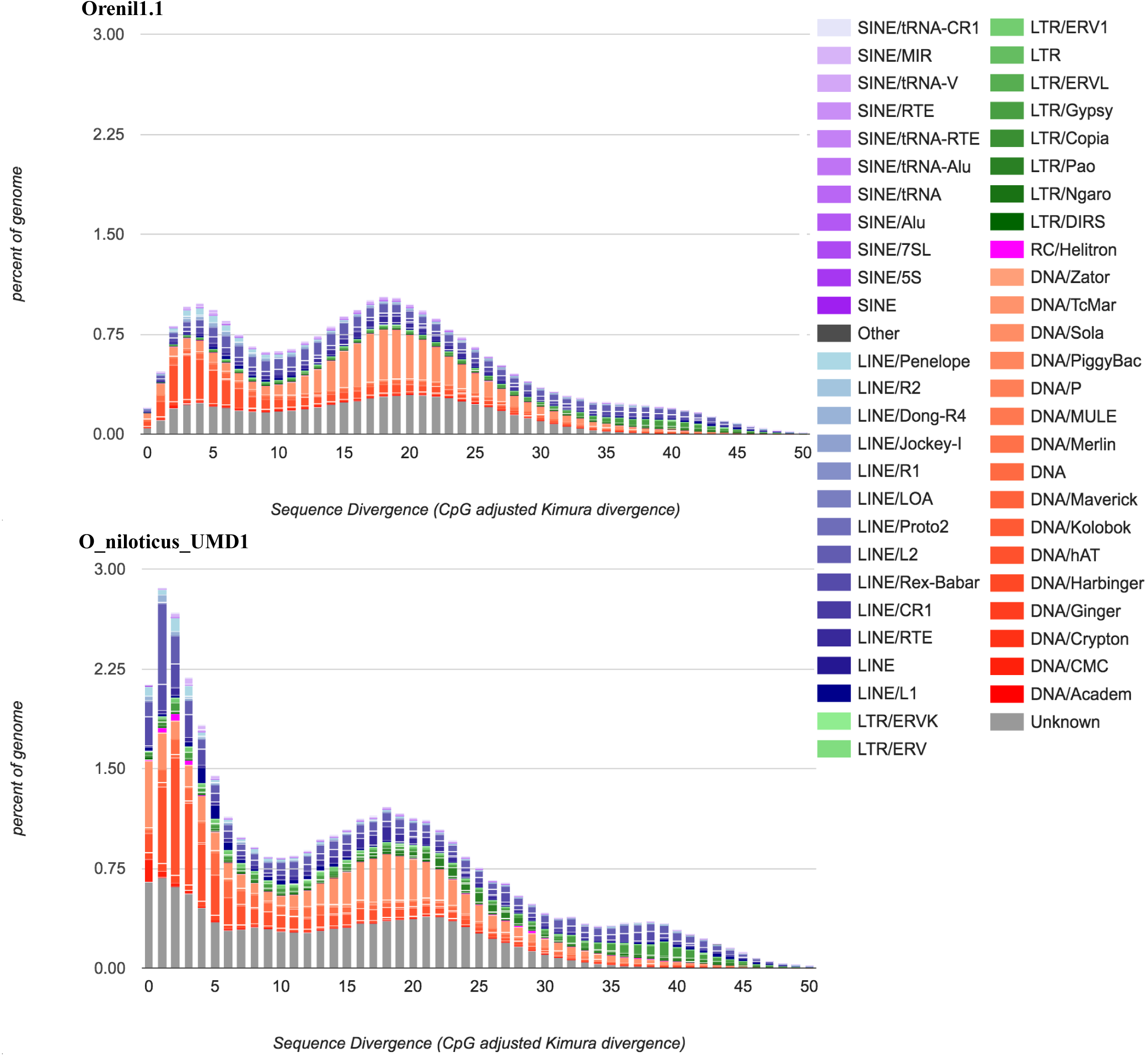
Repeat Landscape comparison. The percentage of both the O_niloticus_UMD1 and Orenil1.1 assemblies that each TE family is represented at in particular substitution levels analogous to the age of TEs (Kimura substitution level – CpG adjusted).

Satellite regions represent one of the most highly repetitive regions of the genome and are often associated with centromeric and heterochromatic regions. Two tilapia-specific satellite repeats have been previously described. ONSATA is a 209bp repeat unit and shows variability between related tilapiine species (38). Only 29 copies of ONSATA (comprising a total of 2,917bp) were assembled and annotated in the original Orenil1.1 assembly. In the new O_niloticus_UMD1 assembly, 226 regions of ONSATA comprising a total of 1,386,985bp were assembled and annotated. Many of the ONSATA regions, the longest of which was 43,805bp on the unanchored contig908, were composed of multiple, nested ONSATA copies. ONSATB is a 1,904bp repeat unit that is organized in tandem arrays and appears to be more conserved and perhaps under selective constraint (39). 48 copies of ONSATB (comprising a total of 11,036bp) were assembled and annotated in the original Orenil1.1 assembly. In the new O_niloticus_UMD1 assembly, 1,481 copies of ONSATB (comprising a total of 2,889,496bp) were assembled and annotated. Again, many of the ONSATB regions were composed of multiple ONSATB copies, the longest of which was 11,210bp located near the beginning of LG12 (607,345-618,555).

TEs specific to African cichlid species have been previously sequenced and used as molecular markers to study evolutionary history and phylogenetics of African cichlids (40,41). Some of these African cichlid specific or “AFC” LINEs and SINEs had been previously assembled and annotated in the Orenil1.1 assembly. An additional 2.3Mbp of AFC-specific TE sequence was annotated in the new O_niloticus_UMD1 assembly. This 2.3Mbp increase was assembled across 55 fewer AFC TE copies, which resulted in longer mean length AFC TE copies. This suggests that the previous assembly contained many fragmented AFC specific TE copies.

### Recently Duplicated Regions

Recently duplicated genes are notoriously difficult to assemble due to their high sequence identity (18). Using short Illumina reads to assemble these regions is a difficult task even with mate-pair sequence data across multiple spatial scales. In a previous study of the tilapia *vasa* gene, we identified three partial gene sequences in the Orenil1.1 assembly (42). We then screened a tilapia BAC library for *vasa* gene sequences and identified three BAC clones containing *vasa* sequences. The three clones came from separate restriction fingerprint contigs (43), and represent duplications of the ancestral *vasa* gene. Sanger sequencing identified a full-length *vasa* gene in each of these BAC clones. Fig 3a shows how the previous Orenil1.1 assembly failed to correctly assemble any of the three *vasa* gene copies. Fig 3b indicates how these genes were assembled from each of the BAC clones. Fig 3c details how the new O_niloticus_UMD1 assembly correctly assembles the three copies of the *vasa* gene corresponding to the three BAC clones.

**Fig 3.**
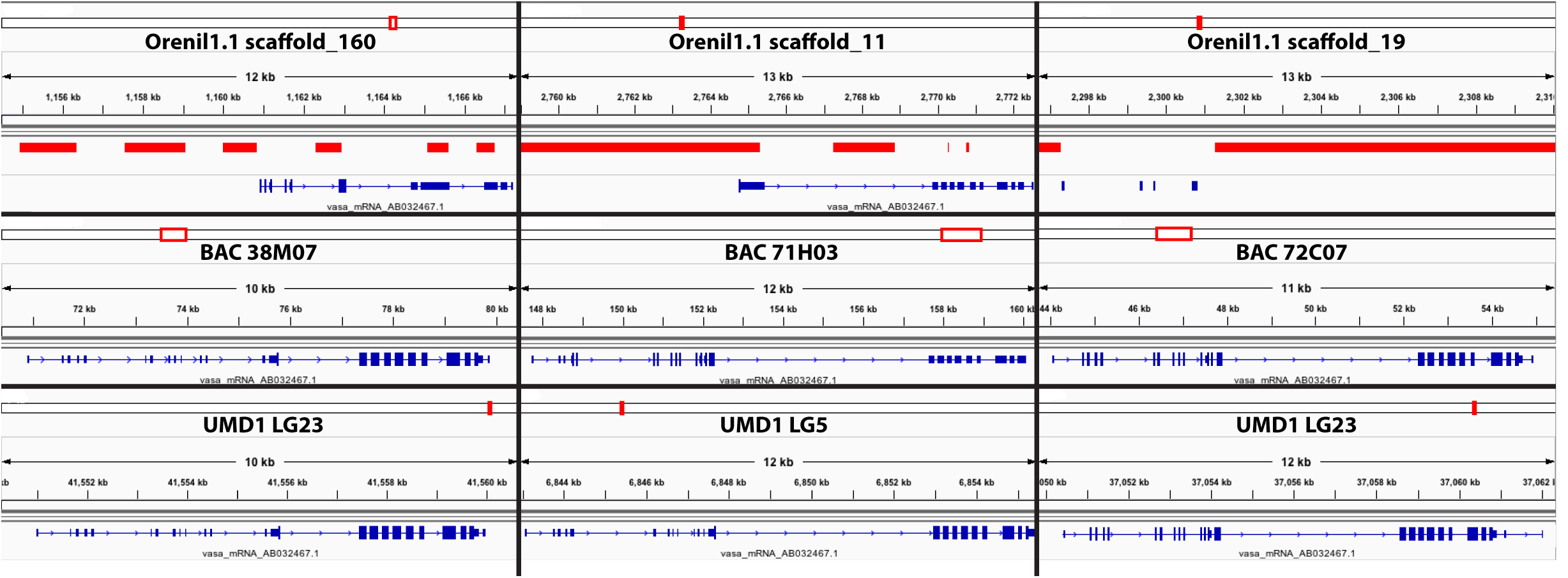
*Vasa* gene duplication. The top row shows the *vasa* transcript (NCBI accession number AB032467.1) aligned to Orenil1.1 assembly scaffolds with gaps shown in solid red. The middle row shows this same *vasa* transcript aligned to the separate BAC assemblies (NCBI accession numbers AB649031-AB649033). The bottom row shows the *vasa* transcript aligned to O_niloticus_UMD1 LGs.

### Sex Determination Regions

The new O_niloticus_UMD1 assembly was used to study sequence differentiation across two sex-determining regions in tilapias. The first region is an XX/XY sex-determination region on LG1 found in many strains of tilapia (9,34,44–47). We previously characterized this region by whole genome Illumina re-sequencing of pooled DNA from males and females (48). We realigned these sequences to the new O_niloticus_UMD1 assembly and searched for variants that were fixed in the XX female pool and polymorphic in the XY male pool. Fig 4 shows the FST and the sex-patterned variant allele frequencies for XX/XY *O. niloticus* comparison across the complete Orenil1.1 and O_niloticus_UMD1 assemblies, while Fig 5 focuses on the highly differentiated ~9Mbp region on LG1 with a substantial number of sex-patterned variants, indicative of a reduction in recombination in a sex determination region that has existed for some time (48).

**Fig 4.**
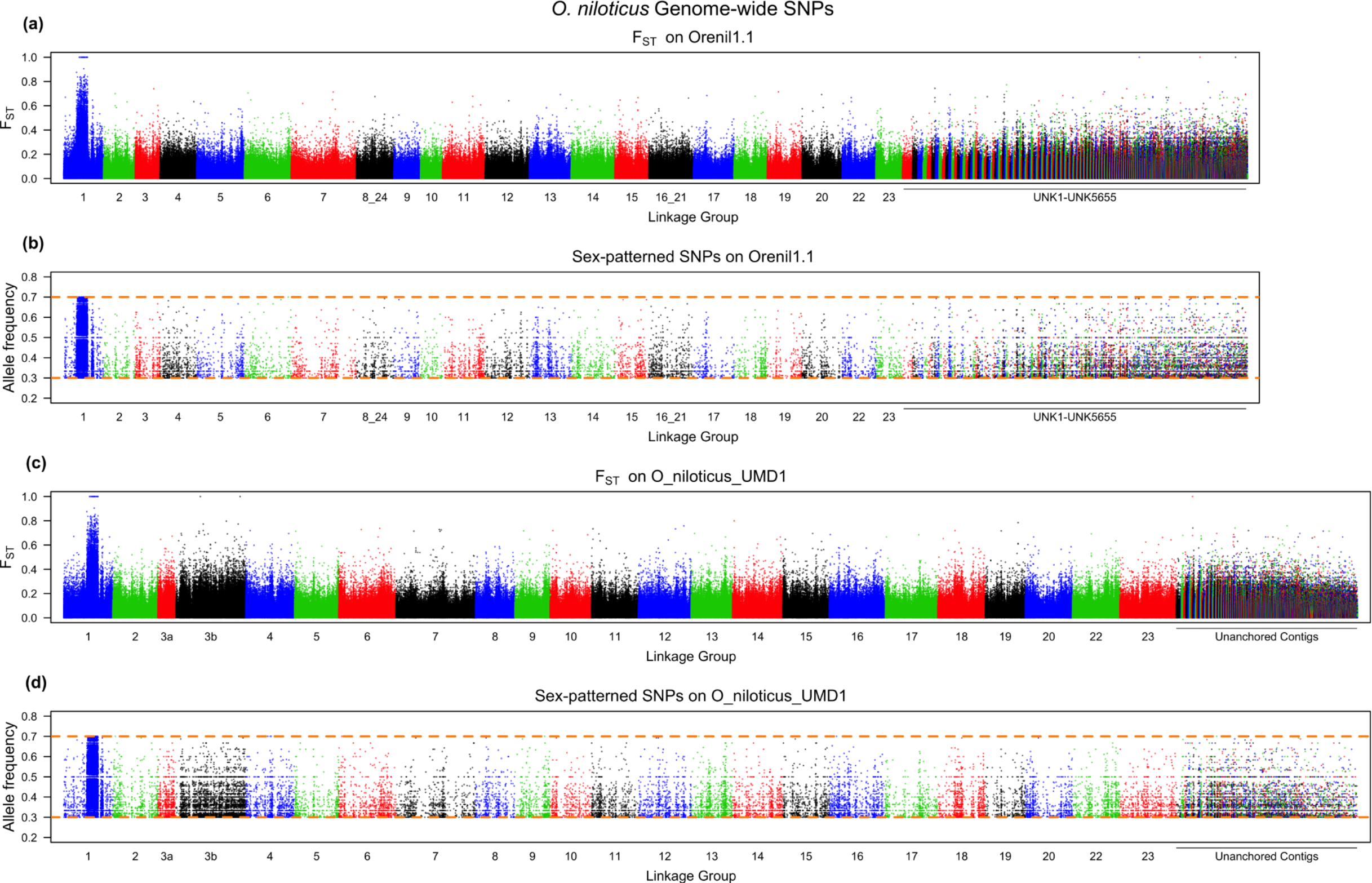
Whole genome *O. niloticus* sex comparison. **a)** F_ST_ comparison of XX female pool versus XY male pool on Orenil1.1. **b)** Sex-patterned variants across Orenil1.1. **c)** F_ST_ comparison of XX female pool versus XY male pool on O_niloticus_UMD1. **d)** Sex-patterned variants across O_niloticus_UMD1.

**Fig 5.**
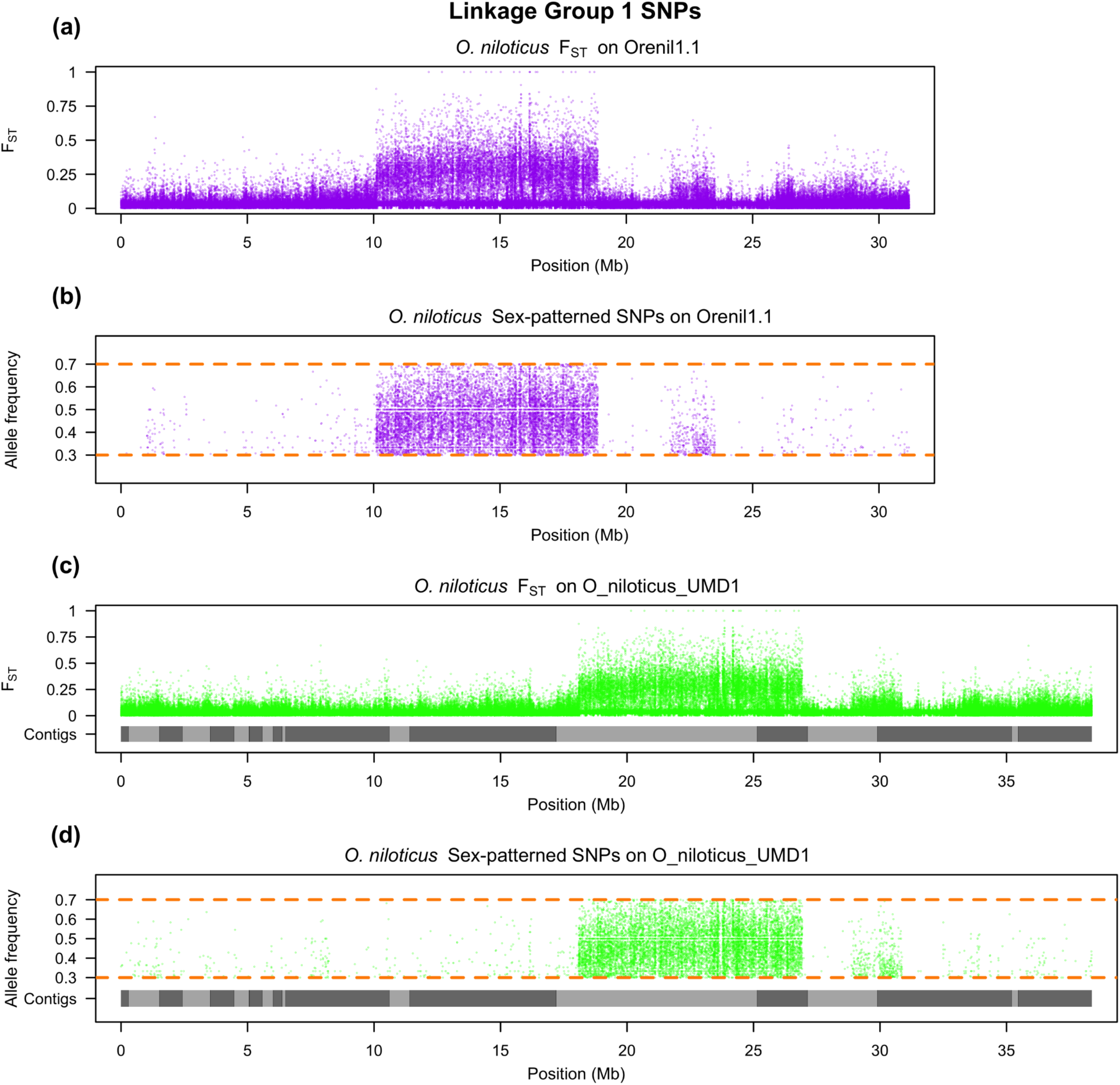
LG1 *O. niloticus* sex comparison. **a)** F_ST_ comparison of XX female pool versus XY male pool on LG1 of Orenil1.1. **b)** Sex-patterned variants on LG1 of Orenil1.1. **c)** F_ST_ comparison of XX female pool versus XY male pool on LG1 of O_niloticus_UMD1. Anchored contig boundaries are depicted with grey bars. **d)** Sex-patterned variants on LG1 of O_niloticus_UMD1.

The second sex comparison is for an ZZ/WZ sex-determination region on LG3 in a strain of *O. aureus* (11,49). This region has not previously been characterized using whole genome sequencing. For this comparison we identified variants alleles fixed in the ZZ male pool and polymorphic in the WZ female pool. Fig 6 shows the F_ST_ and the sex-patterned variant allele frequencies for this comparison across the whole O_niloticus_UMD1 assembly, while Fig 7 focuses on the differentiated region on LG3. *O. aureus* LG3 contains a large ~50Mbp region of differentiated sex-patterned variants, also indicative of a reduction in recombination in the sex determination region. Fig 6 also shows this differentiation pattern on several other LGs (LG7, LG9, LG14, LG16, LG18, LG22 and LG23). It is possible that these smaller regions of sex-patterned differentiation are actually translocations in *O. aureus* relative to the *O. niloticus* genome assembly.

**Fig 6.**
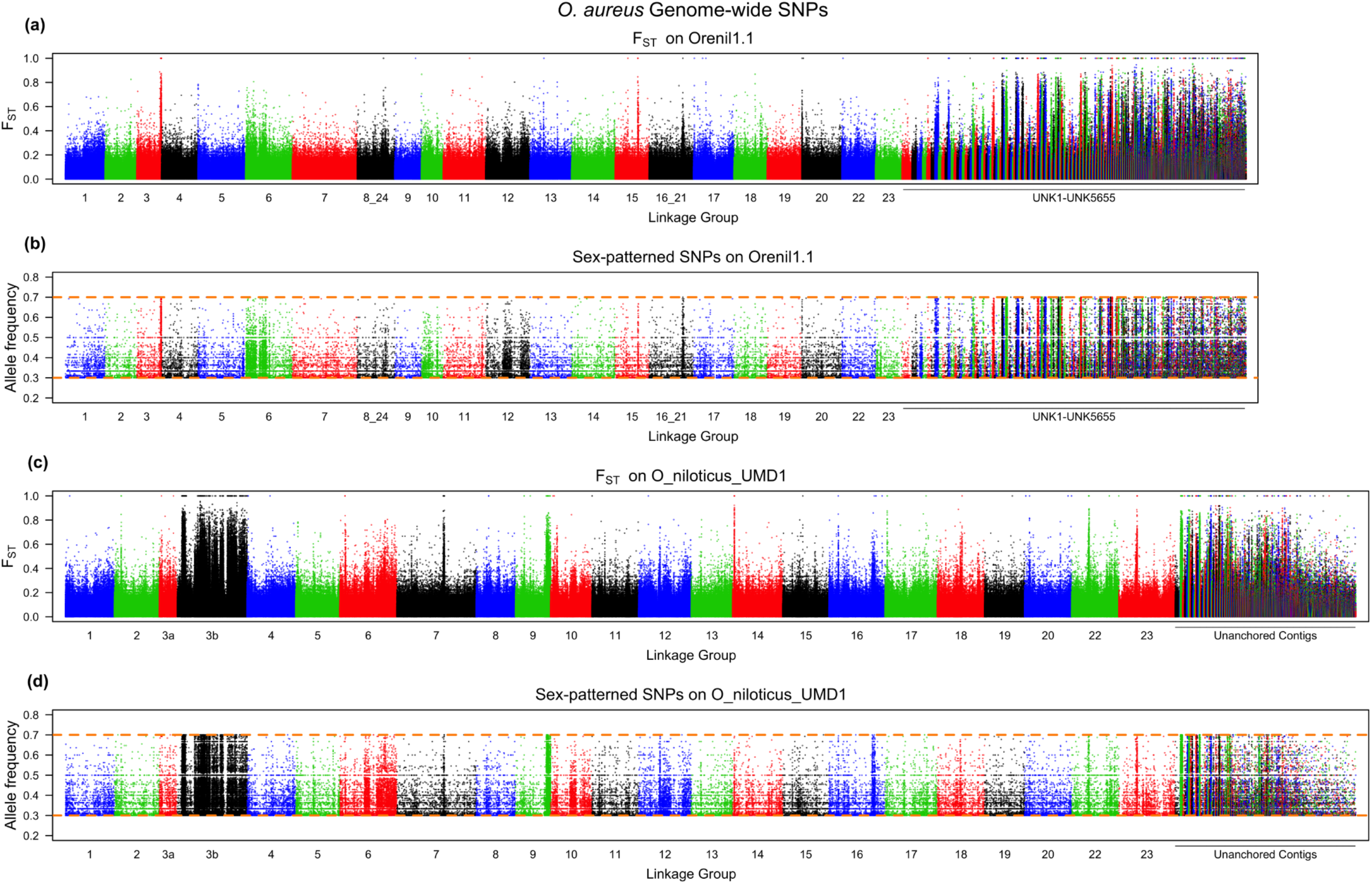
Whole genome *O. aureus* sex comparison. **a)** F_ST_ comparison of ZW female pool versus ZZ male pool on Orenil1.1. **b)** Sex-patterned variants across Orenil1.1. **c)** F_ST_ comparison of ZW female pool versus ZZ male pool on O_niloticus_UMD1. **d)** Sex-patterned variants across O_niloticus_UMD1.

**Fig 7.**
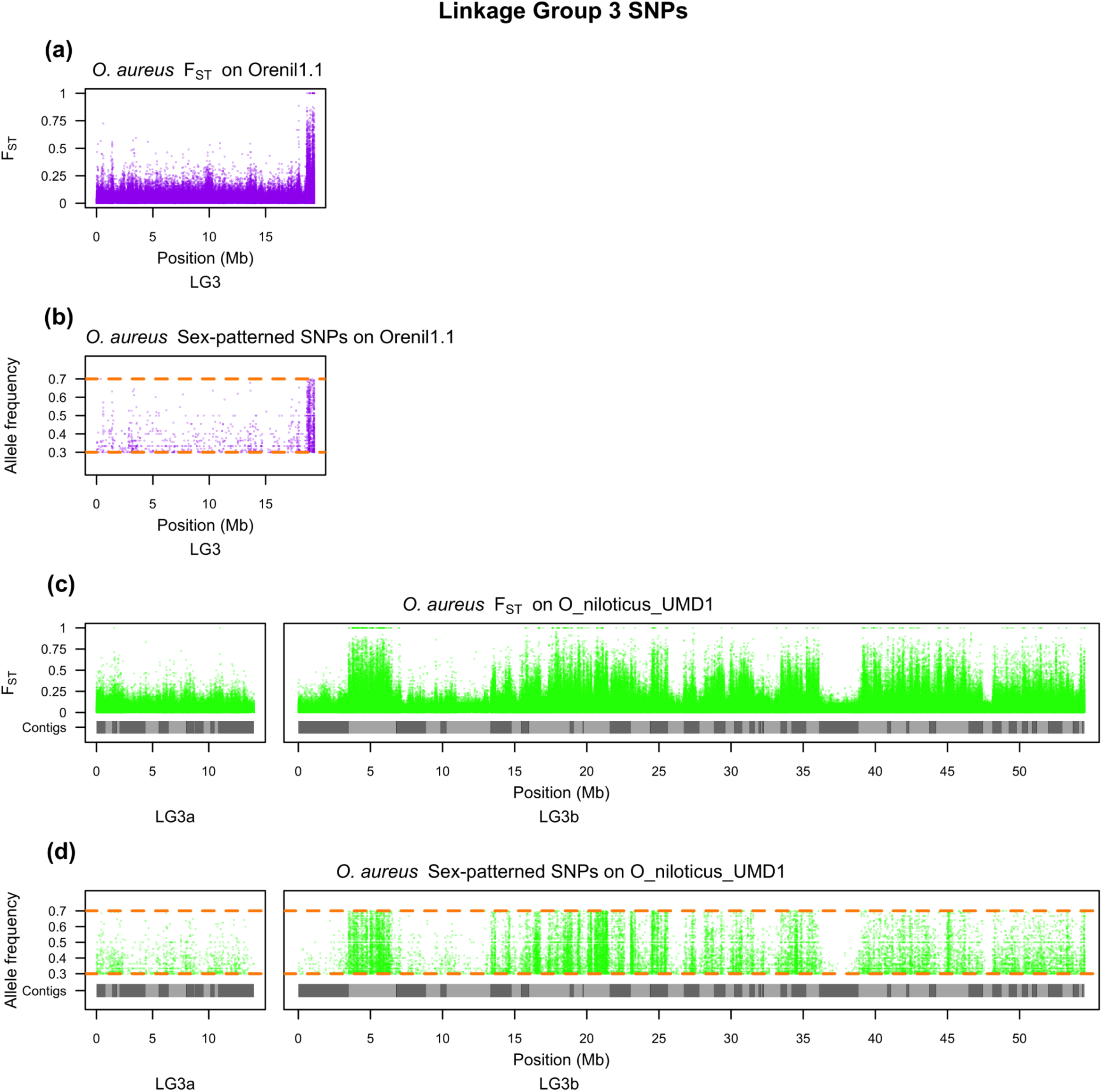
LG3 *O. aureus* sex comparison. **a)** F_ST_ comparison of ZW female pool versus ZZ male pool on LG3 of Orenil1.1. **b)** Sex-patterned variants on LG3 of Orenil1.1. **c)** F_ST_ comparison of ZW female pool versus ZZ male pool on LG3 of O_niloticus_UMD1. Anchored contig boundaries are depicted with grey bars. **d)** Sex-patterned variants on LG3 of O_niloticus_UMD1.

The overall number of sex-patterned variants was markedly increased for both sex comparisons using the new assembly. Table 8 indicates this and provides the number of sex-patterned variants in each comparison across the whole genome as well as on the respective sex-determination LG. LG3 saw the largest gain of sex-patterned variants (1,445 to 24,983 variants) due to the fact that the LG3 assembly now includes 49.3Mbp of new sequence (Table 3).

**Table 8.** LG1 and LG3 sex-patterned variants using both assemblies.

|  | Orenil1.1 | O_niloticus_UMD1 |
| --- | --- | --- |
| ***O. niloticus*** |  |  |
| LG1 sex-patterned variants | 11,894 | 12,225 |
| Non-LG1 sex-patterned variants | 17,579 | 26,493 |
| Total sex-patterned variants | 29,473 | 38,718 |
| ***O. aureus*** |  |  |
| LG3 sex-patterned variants | 1,445 | 24,983 |
| Non-LG3 sex-patterned variants | 79,936 | 78,423 |
| Total sex-patterned variants | 81,381 | 103,406 |

## Discussion

### Genome Assembly

We explored the parameter space of both the FALCON and Canu genome assembly packages and produced 37 candidate assemblies (Table S1). Since the true sequence is not known, we had to deduce which of the candidate assemblies best represented the true sequence of the homozygous clone. We elected to assess the assemblies with a variety of metrics, and to select the assembly that scored well across all of the most important metrics.

The first metric is the overall size of the assembly, which should closely match the estimated size of the genome. The size of the *O. niloticus* genome has been measured by both Feulgen densitometry and bulk fluorometric assay. Five separate measurements range between 0.95-1.20 picograms or ~0.929-1.174Gbp (35). The average genome size of these five estimates is 1.082Gbp. The various assemblies ranged in size from 975.1Mbp to 1.07Gbp. The assembly that was chosen (#14) has a length of 1.01Gbp, which corresponds to 93.3% of the estimated size of the genome.

The second set of metrics we considered were the standard measures of assembly contiguity such as NG50, number of contigs, longest contig and mean contig size. The third set of metrics consisted of assembly likelihood (ALE) scores, which were calculated by aligning four Illumina libraries (fragment, 3kbp, 6-7kbp, and 40kbp – Table 9, Methods) as well as the 44X PacBio library against each candidate assembly. The fourth metric measured the accuracy of the assemblies at larger scales by aligning the contigs to a ~29X clone coverage library of ~150kbp BAC-end sequences (32) and to existing genetic and physical maps of *O. niloticus* (33,34). Alignment of the RH and RAD maps to the candidate assemblies indicated that every assembly had a relatively low and consistent number of misassemblies (Table S1). The fifth metric assessed the completeness of each candidate genome assembly by looking for two core eukaryotic gene sets, CEGMA (29) and BUSCO (30).

No candidate assembly ranked the best for all of these different metrics. In order to choose a preferred assembly, we used principal component analysis to organize the several scores for each assembly. The PCA analysis showed a noticeable difference between the Canu assemblies and the FALCON assemblies (S1 Fig). All of the Canu assemblies clustered together in PCA space. The FALCON assemblies fell into two separate clusters because five of the FALCON assemblies (#17, 32, 34, 35, and 36, Table S1) had low ALE scores and NG50s. The other FALCON assemblies tended to show overall better ALE scores for the 44X PacBio library than did the Canu assemblies. This is due to differences in the consensus accuracy between Canu and FALCON assemblies. The 44X PacBio ALE placement and insert scores were virtually the same across all candidate assemblies, but the 44X PacBio ALE k-mer scores were lower for the Canu assemblies. This suggests a slight difference in consensus between Canu and FALCON, although it is probably not noticeable after the polishing steps.

Leaving aside the five low quality FALCON assemblies, a major tradeoff in the PCA is between size of the assembly and the PacBio ALE score. The FALCON assemblies are all smaller than the Canu assemblies, and for the reasons discussed above, have higher ALE scores for the PacBio library. We elected to focus on the Canu assemblies, where the major tradeoff is between the quality of the assembly (ALE scores, NG50, completeness) and size of the assembly (Total size, exon bp mapped). Ultimately, we chose the assembly (Canu #14) with the best overall ALE average rank. This assembly was 28.8Mbp shorter than the longest Canu assembly (#15).

Alignment of the RH and RAD maps to the candidate assemblies indicated that every assembly had a relatively similar and low number of misassemblies (Table S1). To correct these misassemblies in the polished version of assembly #14, the locations of misassemblies were first narrowed using the RH and RAD map data together. This typically narrowed the location of a misassembly to a region of less than 1Mbp. From there, the region around each misassembly breakpoint was inspected using alignments of the PacBio data, Illumina data, RefSeq gene set, BAC-end sequences as well as the RepeatMasker annotations. A characteristic signal of high variation in the PacBio alignments, low physical coverage in the Illumina libraries (best characterized with the largest 40kbp Illumina library), and a high density of large and nested repeats was seen in each region of misassembly. Regions of high variation in the PacBio alignments and low 40kbp physical coverage were then calculated genome-wide to investigate whether additional misassemblies might be hidden in the assembly. When considering the PacBio highly variant regions and the low physical 40kbp coverage regions individually, both sets over-estimated the number of misassembly regions. These false-positive potential misassemblies occurred in regions where there was support for correct and continuous assembly based on both RH and RAD map alignments, which together lend stronger support. Only in two cases were there regions that had high PacBio variation, low physical 40kbp coverage and no alignment of RH or RAD map data. We decided to break the assembly at these two locations as well.

### Anchoring

A total of 868.6Mbp of the assembled contigs were anchored to the 22 LGs in O_niloticus_UMD1. Overall, 258Mbp of additional (non-gap) sequence has been anchored in the O_niloticus_UMD1 assembly (Table 3). All but two of the O_niloticus_UMD1 LGs (LG5 and LG13) are larger in size than in the previous Orenil1.1 assembly. LG5 is 2.7Mbp smaller and LG13 is 0.4Mbp smaller. It is possible that the Orenil1.1 assembly correctly assembled more of these LG5 and LG13. Alternatively, the size difference could be due to overestimates of gap sizes in the Orenil1.1 assembly and/or incorrect assignment of contigs/scaffolds to the wrong LG, which have now been correctly assigned.

It should be noted that although two markers were required to anchor and orient any contig to a particular LG, not all of the markers in the RAD map were located at distinct map positions (i.e. the map has multiple markers at the same genetic position). Therefore, in some cases (particularly involving many of the smaller and repetitive contigs that were anchored to LG3b), the orientation of contigs on LGs is ambiguous. We chose to allow anchoring of these contigs to maximize the anchoring of the many small repetitive contigs that make up LG3.

### Annotation

Table 5 provides the RefSeq annotation summary of both the Orenil1.1 and new O_niloticus_UMD1 assemblies. The increase in gene and pseudogene annotations is due to the fact that the O_niloticus_UMD1 assembly contains an additional 189.5Mbp of sequence that was not present in Orenil1.1. These additional annotations include protein-coding genes (2,920, 11.1% increase), non-coding RNAs (5,091, 145.1% increase) and pseudogenes (227, 67.4% increase). At the same time, there was a decrease in the number of partial mRNA (2,657, 87.1% decrease) and partial CDS (2,066, 83.7% decrease) annotations. This is most likely due to the fact that O_niloticus_UMD1 gene annotations are not disrupted by assembly gaps. The remaining partial annotations may represent recent pseudogenes that the annotation pipeline has little way of differentiating.

The NCBI RefSeq annotation pipeline corrects CDS annotations that have premature stop-codons, frameshifts and internal gaps that would disrupt protein-coding sequence. The RefSeq annotation pipeline corrected 743 CDSs in O_niloticus_UMD1 compared to 817 previously for Orenil1.1. These remaining 743 CDS annotations that required corrections may be due to incomplete polishing in the final O_niloticus_UMD1 assembly, but this number is less than the amount of corrected CDSs annotated in the smaller Orenil1.1 assembly.

### Repeats

The vast majority of TE families are represented by more sequence in the new assembly (Table 7 and Table S2). It is likely that the fragmented Orenil1.1 assembly caused there to be an inflated count of annotated TE copies in places where gaps were inserted within TE copies. The O_niloticus_UMD1 assembly has assembled TE families in longer overall copies than in Orenil1.1 It is also likely that having longer repeat copies and overall 146Mbp more repeat sequence allowed for more accurate annotation of all repeat sequences. In turn, several TE families (such as SINE tRNA-V and LINE Dong-R4, Table S2) have decreased in overall number in the O_niloticus_UMD1 assembly, which is likely due to these TEs being more accurately annotated as different, but related TEs. The most recent and less diverged TE copies have been assembled in far greater number in the new O_niloticus_UMD1 assembly (Fig 2).

The two tilapia-specific satellite repeats, ONSATA (38) and ONSATB (39), have been shown to be present in high copy number. Both of these satellite repeats have previously been physically mapped using fluorescent *in situ* hybridization (FISH) in *O. niloticus* (50). ONSATA was found almost solely in the centromeres, while ONSATB was also scattered throughout the length of each chromosome arm. Consistent with this, we found nested ONSATA repeat segments assembled near the very ends of several anchored chromosomes (LG3b, LG4, LG8, LG14, and LG17). ONSATB nested repeat segments were found near one or both ends of several anchored chromosomes (LG2, LG3a, LG3b, LG4, LG6, LG11, LG12, LG14, LG16, LG17, LG18, LG19, LG20, and LG23). These data suggest that our assembly of these chromosomes extend into the centromeres. These satellite nested repeats were also abundant in several of the misassembled regions (Table 2) suggesting that they remain an obstacle to complete assembly of the genome.

### Recently Duplicated Regions

As the recent *vasa* gene duplication in *O. niloticus* (Fig 3) shows, the use of long reads has enabled the assembly of such recently duplicated regions. It is likely that there are many other recently duplicated regions that have now been assembled. This is supported by the genome completeness analysis with BUSCO that showed there were 26 additional duplicated BUSCOs out of 3023 searched (Table 4). Even though this is a small percentage of the genes analyzed (0.86%), when extrapolated over all the genes in the genome this would amount to hundreds of recently duplicated genes being assembled for the first time. The RefSeq annotation shows that the O_niloticus_UMD1 assembly contained 227 additional pseudogenes (67.4% increase from the Orenil1.1 assembly), which also supports this notion.

### Sex Determination Regions

Manipulation of sex-determination in tilapia has important economic implications. The O_niloticus_UMD1 assembly was used to confirm the known and previously described *O. niloticus* ~9Mbp sex-determination region on LG1 (48). The size and pattern of sex differentiation on LG1 and across the genome is similar in both the Orenil1.1 and O_niloticus_UMD1 assemblies (Fig 4 and Fig 5). A total of 331 additional LG1 sex-patterned variants are identified in the O_niloticus_UMD1 assembly.

The sex-determination region in *O. aureus* is located on the large and highly repetitive LG3. Due to the fact that LG3 is highly repetitive, it was poorly assembled in Orenil1.1 and the vast amount of sex-patterned variants were previously found on unanchored contigs and scaffolds (Fig 6a and 6b). An additional 23,538 LG3-specific *O. aureus* sex-patterned variants are identified in the O_niloticus_UMD1 assembly. Now that LG3 has been assembled and anchored into a much larger LG (68.5Mbp versus 19.3Mbp, Table 3), many of these sex-patterned variants are confirmed on LG3 (Fig 6c and 6d). There still exist a substantial number of sex-patterned variants on unanchored contigs in the new assembly. The overall pattern of *O. aureus* sex differentiation on LG3 is characterized by several sharp transitions between low and high differentiation (e.g. ~5Mbp and ~37Mbp, Fig 7c and 7d). These sharp transitions may be explained by either errors in the anchoring process or structural differences between the reference species (*O. niloticus*) and *O. aureus*. Indeed, there are several peaks of differentiation on other LGs (LG7, LG9, LG14, LG16, LG18, LG22 and LG23, Fig 6). These may also be chromosomal translocation differences between the two species that will need to be investigated further with FISH.

## Conclusions

This study provides a new assembly and annotation of the Nile tilapia *O. niloticus* (O_niloticus_UMD1), which provides a high-quality reference for the cichlid research community as well as one for studying the evolution of vertebrate genomes. The study also serves as a template for vertebrate genome assembly with current technology and describes many of genomic features that can now be represented correctly. Generation of O_niloticus_UMD1 began by comparing candidate *de novo* assemblies systematically comparing them to select a single best assembly. A small number of misassemblies present in this candidate assembly were identified using several different datasets and subsequently corrected. The final anchored O_niloticus_UMD1 assembly remained very contiguous with a contig NG50 of 3.1Mbp and 86% of contigs anchored to LGs. The number of annotated genes increased 27.3% from the previous assembly of *O. niloticus.* Additionally, a vast amount of repetitive sequences (~146Mbp) were added in the O_niloticus_UMD1 assembly, many of which represent very recent TEs. Finally, the O_niloticus_UMD1 assembly was used to better characterize two large sex-determination regions. The first is a ~9MBp region in *O. niloticus* and the second is a ~50Mbp region in the related species *O. aureus*. Further characterization of these sex-determination regions will have important economic implications for farmed tilapia.

## Materials and Methods

### PacBio Sequencing

PacBio sequencing was performed on a new individual from the same XX homozygous clonal line used for the previous whole genome sequencing of *O. niloticus* (16). This mitogynogenetic line was developed and maintained at the University of Stirling, UK (26). All working procedures complied with the UK Animals (Scientific Procedures) Act (51).

The Qiagen MagAttract HMW DNA kit was used to extract high-molecular weight DNA from a nucleated blood cell sample of the female “F11D_XX” individual. Size selection was performed at the Genomics Resource Center, Institute for Genome Sciences using a Blue Pippin pulse-field gel electrophoresis instrument. A library was constructed and 63 SMRT cells were sequenced on their PacBio RS II instrument using the P6-C4 chemistry.

### Assembly

Both Canu (27) (*version 1.0*) and FALCON (22) (*versions 0.3.0 and 0.5.0*) were run to generate candidate *de novo* genome assemblies. The wide range of parameters tested for both algorithms are provided in Table S3. The final assembly (#14), chosen based on the evaluation and likelihood calculations (see below), was run using Canu with the following relevant parameters: ‘*minReadLength=7000 minOverlapLength=2000 MhapSensitivity=high genomeSize=1g errorRate=0.025 - pacbio-raw*’.

### Assembly Accuracy Measurements

Assembly summary metrics were calculated using the assemblathon_stats.pl script (52). Illumina libraries generated previously (16) were aligned to each candidate *de novo* assembly using Bowtie2 (*version 2.2.5* in ‘--*very-sensitive*’ mode). The four different insert size Illumina libraries used are presented in Table 9.

**Table 9.** *O. niloticus* Illumina libraries used for ALE calculations and Pilon polishing.

| Insert size | NCBI SRA accession IDs | Combined coverage | Platform(s) |
| --- | --- | --- | --- |
| Fragment | SRR071589, SRR071593, SRR071594, SRR071601, SRR071604, SRR071605, SRR071610, SRR071619 | 51.6x | Illumina Genome Analyzer II |
| 3kbp | SRR071588, SRR071591, SRR071597, SRR071599, SRR071603, SRR071612, SRR071614 | 196.3x | Illumina HiSeq 2000 |
| 6-7kbp | SRR071602, SRR071607, SRR071615, SRR071616, SRR071617, SRR071620, SRR071622, | 24.6x | Illumina Genome Analyzer II |
| 40kbp | SRR071595, SRR071598, SRR071611 | 4.8x | Illumina Genome Analyzer II |

For each SRA run, raw reads were downloaded from NCBI using the ‘*fastq-dump*’ program from the SRA Toolkit (53) (*version 2.5.2*). Raw fastq files were combined for each insert size group and Trimmomatic (54) (*version 0.32*) was run on the combined fastq files. The 101bp fragment library and 3kbp library reads were each trimmed with the following Trimmomatic settings:

*‘ILLUMINACLIP:TruSeq2-PE.fa:2:30:10 SLIDINGWINDOW:4:20 LEADING:10 TRAILING:10 CROP:101 HEADCROP:0 MINLEN:80’*

The 36bp 6-7kbp library reads were trimmed with the following settings:

*‘ILLUMINACLIP:TruSeq2-PE.fa:2:30:10 SLIDINGWINDOW:4:20 LEADING:10 TRAILING:10 CROP:36 HEADCROP:0 MINLEN:31’*

The 76bp 40kbp library reads were trimmed with the following settings:

*‘ILLUMINACLIP:TruSeq2-PE.fa:1:30:10 SLIDINGWINDOW:10:20 LEADING:5 TRAILING:10 CROP:76 HEADCROP:4 MINLEN:70’*

For the fragment library, the trimmed and filtered reads were next overlapped with FLASH (55) (*version 1.2.11*) using the following parameters: ‘-*m 20-x 0.15-z*’.

The samtools (56) (*version 1.1*) ‘*view*’ and ‘*sort*’ commands were used to convert the Bowtie2 SAM outputs to BAM format. The Picard (57) (*version 2.1.0*) ‘*MarkDuplicates*’ program was run on each of these Bowtie2 alignments with ‘*REMOVE_DUPLICATES=true*’.

The Assembly Likelihood Estimator (ALE) (28) was then run on each of these filtered BAM files to generate likelihood statistics for each candidate Canu and FALCON *de novo* assembly for each Illumina library. Additionally, to generate ALE scores for the raw PacBio data aligned to each assembly, the 44X raw PacBio reads were aligned using BLASR (58) (version *1.3.1.127046*) with the following parameters: ‘-*minMatch 8 -minPctIdentity 70 -bestn 1 -nCandidates 10 -maxScore - 500 -nproc 40 -noSplitSubreads -sam’.* ALE was then run on these BLASR alignments as well.

A set of *O. niloticus* paired BAC-end sequences (32) were aligned against each candidate assembly using BLAST (59–61) (*version 2.3.0+*). The top hit with an *E-value* less than 1e-150 were kept and then assigned a category of alignment relative to the candidate assemblies according to the details described previously (32) and briefly explained for Table S1.

To evaluate the completeness of the candidate assemblies, BUSCO (30) (*version 1.22*) was run (in ‘-*m genome*’ mode) using the ‘*vertebrata*’ lineage-specific profile library. CEGMA (29) (*version 2.5*) was also run on each of the candidate assemblies. CEGMA was run optimized for vertebrate genomes (option ‘--*vrt*’) and relied on GeneWise (*version 2.4.1)*, HMMER (*version 3.1b1*), and NCBI BLAST+ (*version 2.3.0+*) using the provided set of 248 CEGs.

### Principal Component Analysis

The following metrics were calculated and culled for each of the 37 candidate assemblies: Total ALE score for the aligned Illumina fragment, 3kbp, 6-7kbp, and 40kbp libraries; Total ALE score for the aligned PacBio library; Total number of complete CEGs as defined by CEGMA; Longest contig; NG50; Total assembly size (bp); Total number of RefSeq exon bp mapped. *O. niloticus* RefSeq transcripts (31) (*release 70*) were aligned to each of the candidate assemblies using GMAP (62) (*version 2015-07-23*) and exon bp mapped were calculated from the output GFF3 file. R version 3.2.3 was used to perform the PCA analysis using the ‘*prcomp*’ function with ‘*center=TRUE, scale=TRUE*’ and to create plots with the ‘*biplot*’ function.

### Polishing the Assembly

SMRT-Analysis (63) (*version 2.3.0.140936*) was used for polishing the Canu #14 assembly using the 44X raw PacBio reads. First, each SMRT cell was separately aligned to the unpolished Canu assembly using pbalign (*version 0.2.0.138342*) with the ‘--*forQuiver*’ flag. Next, cmph5tools.py (*version 0.8.0*) was used to merge and sort (with the ‘--*deep*’ flag) the pbalign .h5 output files for each SMRT cell. Finally, Quiver (*GenomicConsensus version 0.9.2* and *ConsensusCore version 0.8.8*) was run on the merged and sorted pbalign output to produce an initial polished assembly.

Pilon (64) (*version 1.18*) was run on the intermediate Quiver-polished assembly produced above. Again, Bowtie2 (*version 2.2.5* in ‘--*very-sensitive*’ mode) was used to align Illumina reads to this intermediate assembly for Pilon polishing. The fragment library alignment was supplied to Pilon with ‘--*unpaired*’ while the other 3 insert library alignments were specified with ‘--*jumps*’. Additionally, Pilon was run with the following parameters: ‘--*changes --vcf --chunksize 40000000 --fix all*’.

### Detecting Misassemblies

The 44X coverage raw PacBio reads were aligned to the Quiver- and Pilon-polished Canu #14 assembly using BLASR (58) (version *1.3.1.127046*) with the same parameters as mentioned above. Variants were called using FreeBayes (65) (version *v1.0.2-33-gdbb6160-dirty*). To facilitate FreeBayes processing, regions of the polished assembly were broken into 500kbp chunks using the FreeBayes “fasta_generate_regions.py” script. The separate VCF output files were then concatenated using the VCFtools (66) ‘*vcf-concat*’ program. The FreeBayes utility ‘*vcffilter*’ was used to filter these variants for quality greater than 10 (‘-*f "QUAL > 10”* ‘). VCFtools was then used to compute variant density by specifying ‘--*SNPdensity 10000*’ to calculate variant density in 10kbp windows. Highly variant regions were flagged if there were more than 1 variant per 1kbp over a 10kbp window.

The 40kbp mate-pair Illumina reads of the same homozygous inbred *O. niloticus* line (16) were downloaded from the NCBI SRA (SRR071595, SRR071598, and SRR071611). Trimmomatic (54) (*version 0.32*) was run to remove adaptor sequences and to trim/quality filter these reads. The relevant parameters for Trimmomatic were ‘*PE -phred33 ILLUMINACLIP:TruSeq2-PE.fa:1:30:10 SLIDINGWINDOW:10:20 LEADING:5 TRAILING:10 CROP:76 HEADCROP:4 MINLEN:70*’. The trimmed and filtered reads were combined and aligned to the polished assembly using BWA mem (67) (*version 0.7.12-r1044*) with the ‘-*M*’ flag. The Picard (57) (*version 2.1.0*) ‘*SortSam*’ program was used to convert the SAM output to BAM (*‘SORT_ORDER=coordinate’*) and the Picard ‘*MarkDuplicates*’ program was used to identify duplicate reads. The physical coverage of the 40kbp mate-pairs was calculated on a per-contig basis using a series of piped samtools (56) (*version 1.1*) and bedtools (*version v2.26.0*) commands using the following template, where ‘*contig*’ and ‘*contig_size*’ are the specific contig and its respective size: ‘*samtools sort -no <(samtools view -bh -F 2 -q 1 40kb.bam contig) tmp | bamToBed -i stdin -bedpe | cut -f 1,2,6 | sort -k 1,1 | bedtools genomecov -i stdin -g <(echo -e "contig\tcontig_size\n") -bga -pc | grep contig > output*’. Regions within 200kbp of the start or end of a contig were then excluded from this analysis. Regions below 20x physical coverage of 40kbp mate-pair reads were flagged.

Regions of high variant density within 20kbp of each other, based on raw PacBio alignments, were merged using the bedtools ‘*merge*’ program (*‘-d 20000*’). The same merging of windows was performed for regions of low physical coverage based on the 40kbp mate-pair library. The bedtools ‘*intersect*’ program was then used to determine regions of high-density PacBio variants and low 40kbp mate-pair physical coverage that overlapped by at least 80% in the high-density PacBio variants merged windows (*‘-f 0.8*’).

Regions of both high-density PacBio variants and low 40kbp mate-pair physical coverage were compared to the alignments of the RH map and RAD map to confirm or contradict the putative misassemblies. Putative misassembled regions were manually inspected using the BLASR and BWA alignments using IGV (68). In addition to these tracks, both RefSeq (31) (*release 70*) *O. niloticus* transcripts aligned to the polished Canu assembly using GMAP (62) (*version 2015-07-23*) and RepeatMasker (69) repeat annotations were considered when defining the exact location of a misassembly. Break locations were chosen so that they did not occur within RefSeq transcripts or within single repeat annotations. The REAPR (70) (*version 1.0.18*) ‘*break*’ program was used to break and fix the polished Canu assembly by providing the determined break locations.

### Anchoring with Chromonomer

Chromonomer (71) (*version 1.05*) was first used to anchor the polished and misassembly-corrected assembly using the RH map for *O. niloticus* (33). This initial anchored assembly was then subsequently anchored again with a RAD map for *O. niloticus* (34). BWA mem (*version 0.7.12-r1044)* was used in both Chromonomer runs to create the input SAM file by aligning respective map marker sequences to the appropriate intermediate assembly. A minimum of two markers were required to anchor a contig to a particular LG. Gaps of 10kbp were placed between anchored contigs using ‘--*join_gap_size 10000*’ in Chromonomer. Several RH linkage groups required manual placement were fixed by replacing their entries in the SAM file used by Chromonomer. The RH map LGs that were not anchored using the RAD map (“LOD4.9-RH10-LG10”, “LOD6.5-RH17-LG15”, and “LOD5.7-RH31-LG3”) were manually placed onto the final LGs by using the additional mapping data provided in the previous publication, ‘Additional file 4. Data S4’ of (33) which integrated FISH mapping of BAC markers and an previous genetic map (72). Three RH LGs also had to be fixed as they contained a number of repetitive markers, which was causing them to be anchored to incorrect linkage groups in the RAD map (“LOD4.5-RH5-LG9,” LOD6.9-RH6-LG5.rev”, and “LOD5.1-RH8-LG13”).

To further evaluate the candidate assemblies described above, the Chromonomer output file ‘*problem_scaffolds.tsv*’ was used to count the number of contigs in each assembly that had multiple markers that mapped to two or more separate linkage groups.

### RefSeq Annotation

The O_niloticus_UMD1 assembly was submitted to NCBI to perform the Eukaryotic Genome Annotation Pipeline (73). This automated pipeline masks the assembly, and aligns existing transcript, protein, RNA-seq, and curated RefSeq sequences to it. Gene prediction based on these alignments is performed and the best gene models are selected among the RefSeq and predicted models which are then made available as the annotation release. The O_niloticus_UMD1 assembly was annotated as “annotation release 103” (74) and the previous “annotation release 102” (75) consisted of the Orenil1.1 annotation. The numbers in Table 5 were extracted from these summaries.

### Repeat Annotation

The annotation of repetitive elements was run on several of the intermediate assemblies as well as the final O_niloticus_UMD1 assembly. For each of these assemblies, RepeatModeler (76) (*version open-1.0.8*) was first used to identify and classify *de novo* repeat families present in each assembly. These *de novo* repeats were then combined (separately for each assembly) with the RepBase-derived RepeatMasker libraries (77). RepeatMasker (69) (*version open-4.0.5*) was then run on each of these assemblies using NCBI BLAST+ (*version 2.3.0+*) as the engine (‘-*encbi*’) and specifying the combined repeat library (‘-*lib*’). The more sensitive slow search mode (‘-*s*’) was used.

### Analysis of Duplicated Vasa Regions

The *vasa* transcript (NCBI accession AB032467.1) was aligned to three assembled BAC clones (NCBI accessions AB649031-AB649033) corresponding to the three copies of *vasa* present in the *O. niloticus* genome (42) using GMAP (62) (*version 2015-07-23*). The *vasa* transcript was also aligned to the scaffolds of the Orenil1.1 assembly and the final anchored O_niloticus_UMD1 assembly. IGV was used to generate images displaying the transcript alignments of the duplicated *vasa* genes.

### Sex Comparisons

Sex comparisons were run on the O_niloticus_UMD1 assembly for two species of tilapia, *O. niloticus* and *O. aureus*. The *O. niloticus* sequence data used in this study was previously described (48). The *O. aureus* individuals used were F1 individuals derived from a stock originally provided by Dr. Gideon Hulata (Institute of Animal Science, Agricultural Research Organization, The Volcani Center, Bet Dagan, Israel) and maintained at University of Maryland. These animal procedures were conducted in accordance with University of Maryland IACUC Protocol #R-10-74. A total of 58 *O. niloticus* XY males, 33 *O. niloticus* XX females, 22 *O. aureus* ZZ males and 22 *O. aureus* WZ females were pooled separately, sheared to ~500bp on a Covaris shearer, and sequenced on an Illumina HiSeq 2000. The reads from each pool were separately mapped to O_niloticus_UMD1 using BWA mem (*v0.7.12*). The alignments were sorted and duplicates were marked with Picard (*v2.1.0*). Alignments were converted into an mpileup file using Samtools (*v0.1.18*) and subsequently into a sync file using Popoolation2 (*v1201*) (78). Estimates of F_ST_ and analyses of sex-patterned variants (SNPs and short deletions that are fixed or nearly fixed in the homogametic sex and in intermediate frequency in the heterogametic sex) were carried out using *Sex_SNP_finder_GA.pl* (https://github.com/Gammerdinger/sex-SNP-finder). For the *O. niloticus* sex comparison, the XX females were set to be the homogametic sex. For the *O. aureus* comparison, the ZZ males were set to be the homogametic sex.

### Sequence availability

Female *O. niloticus* pool: SRR1606304

Male *O. niloticus* pool: SRR1606298

Female *O. aureus* pool: SRR5121055

Male *O. aureus* pool: SRR5121056

44X *O. niloticus* PacBio reads: SRP093160

O_niloticus_UMD1 Assembly: MKQE00000000

## Acknowledgements

We thank Luke Tallon and Naomi Sengamalay at the Genomics Resource Center, Institute for Genome Sciences for providing a high quality PacBio library and sequence reads. We thank Gideon Hulata of the Agricultural Research Organization in Israel for providing the original stocks of *O. aureus* that we maintained for this study. We thank Christopher Hill at University of Washington for helpful guidance on assembly parameters. We thank the members of the Kocher and Carleton labs at UMD for providing thoughtful advice during the course of the project. We thank Karen Carleton for reading and providing thoughtful feedback on the manuscript. We thank the NCBI RefSeq curators for annotation of the O_niloticus_UMD1 genome assembly. We acknowledge the University of Maryland supercomputing resources (www.it.umd.edu/hpcc/) made available in conducting the research reported in this paper.

### Author Contributions

MAC and TDK conceived and designed study. DJP provided the blood sample from the homozygous clonal line of *O. niloticus* and KLB optimized and performed the HMW DNA extraction. MAC performed the genome assembly, evaluation and analyses. WJG and TDK raised specimens, performed DNA extraction and prepared sequencing library construction of the *O. aureus* samples. WJG organized map data and performed the computational analysis of the sex comparisons. MAC and TDK wrote the paper. All authors read and approved the manuscript.

## Supporting Information

**S1 Fig. PCA analysis of candidate assemblies.** PCA analysis of the 37 candidate assemblies (assembly numbers listed in Table S1 and Table S3) is composed of total size (bp), exon bp mapped, complete CEGMA CEGs, NG50, longest contig, and overall ALE scores (Illumina fragment, 3kbp, 6-7kbp, 40kbp and 44X PacBio libraries).

**S2 Fig. Example misassembly signature.** An example misassembly identified by both RH and RAD maps showing the characteristic signature of high variation in the 44X PacBio read alignments as well as low coverage in the 40kbp Illumina mate-pair library and high density of repetitive elements.

**S1 Table. Extended assembly metrics of the 37 candidate assemblies.** Assembly statistics for Canu assemblies (#1-16), FALCON assemblies (#17-37), total and composite ALE scores for each assembly with rankings, and CEGMA/BUSCO results for each assembly. BAC-end types are defined as: type 1 BAC-ends have only a single read-pair aligned, type 2 BAC-ends have both read-pairs aligned in correct orientation, type 3 BAC-ends have incorrect orientation or mapping distance, and type 4 BAC-end read-pairs match separate contigs.

**S2 Table. Extended repeat annotations of O_niloticus_UMD1 and Orenil1.1.** RepeatMasker results for each TE family and other repeats for both assemblies and the differences between the two.

**S3 Table. Candidate assembly parameters.** Parameters used for the 37 candidate assemblies, Canu and FALCON assembly parameters are separated.

